# Heterogeneity of *dN/dS* ratios at the classical HLA class I genes over divergence time and across the allelic phylogeny

**DOI:** 10.1101/008342

**Authors:** Bárbara Domingues Bitarello, Rodrigo dos Santos Francisco, Diogo Meyer

**Keywords:** balancing selection, HLA, MHC, *dN/dS*, allelic lineages, antigen recognition site, divergent allele advantage

## Abstract

The classical class I HLA loci of humans show an excess of nonsynonymous with respect to synonymous substitutions at codons of the antigen recognition site (ARS), a hallmark of adaptive evolution. Additionally, high polymporphism, linkage disequilibrium and disease associations suggest that one or more balancing selection regimes have acted upon these genes. However, several questions about these selective regimes remain open. First, it is unclear if stronger evidence for selection on deep timescales is due to changes in the intensity of selection over time or to a lack of power of most methods to detect selection on recent timescales. Another question concerns the functional entities which define the selected phenotype. While most analysis focus on selection acting on individual alleles, it is also plausible that phylogenetically defined groups of alleles (“lineages”) are targets of selection. To address these questions we analyzed how *dN/dS* (*ω*) varies with respect to divergence times between alleles and phylogenetic placement (position of branches). We find that *ω* for ARS codons of class I HLA genes increases with divergence time and is higher for inter-lineage branches. Throughout our analyses, we used non-selected codons to control for possible effects of inflation of *ω* associated to intra-specific analysis, and showed that our results are not artifactual. Our findings indicate the importance of considering the timescale effect when analysing *ω* over a wide spectrum of divergences. Finally, our results support the divergent allele advantage model, whereby heterozygotes with more divergent alleles have higher fitness than those carrying similar alleles.

## 1 Introduction

MHC class I and II classical molecules are cell-surface glycoproteins which mediate presentation of peptides to T-cell receptors, and play a key role in triggering adaptive immune responses when the bound peptide is recognized as foreign (Klein and Sato 2000). In humans, they are coded by HLA class I (*HLA-A, -B,* and *-C*) and II (*HLA-DR, -DQ,* and *-DP*) classical genes. The class I and class II HLA classical genes are the most polymorphic in the human genome (Meyer and Thomson 2001), and knowledge about their function in the immune response supports a role for balancing selection in driving the diversity patterns at these loci.

A number of findings suggest MHC genes have experienced balancing selection: unusually high level of heterozygosity with respect to neutral expectations (Hedrick and Thomson 1983); existence of trans-species polymorphisms (Takahata and Nei 1990); high levels of linkage disequilibrium (Huttley et al 1999); site frequency spectra with excess of common variants (Garrigan and Hedrick 2003); high levels of identity-by-descent compared to genomic averages (Albrechtsen et al 2010); positive correlation between HLA polymorphism and pathogen diversity (Prugnolle et al 2005), and significant associations of HLA alleles with the course of infectious diseases (e.g. Apps et al 2013). Information on the crystal structure of MHC molecules (Bjorkman et al 1987) allowed the identification of a specific set of amino acids that make up the antigen recognition site (ARS), which determines the peptides that the molecule is able to bind (Bjorkman et al 1987; Chelvanayagam 1996). The codons of the ARS were shown to have increased nonsynonymous substitution rates (Hughes and Nei 1988, 1989), consistent with the hypothesis that adaptive evolution at HLA loci is driven by peptide binding properties.

Several models of selection are compatible with balancing selection at MHC genes. Heterozygote advantage assumes that heterozygotes have higher fitness values because they are able to mount an immune response to a greater array of pathogens, an idea originally proposed by Doherty and Zinkernagel (1975), who showed that mice which were heterozygous for the MHC had increased immunological surveillance. Heterozygote advantage has received support from experiments in semi-natural populations of mice (Penn et al 2002), which show increased resistance of heterozygotes to multiple-strain infection, and through the finding that among humans infected with HIV, those which are heterozygous for HLA genes have slower progression to AIDS (reviewed in Dean et al 2002). Heterozygote advantage has also received support from substitution rate studies (Hughes and Nei 1988, 1989) as well as simulation-based studies (e.g. Takahata and Nei 1990). A second model for balancing selection at MHC genes is negative frequency dependent selection (or apostatic selection), according to which rare variants have a selective advantage over common ones, because pathogens are more likely to evade presentation by common molecules (Slade and McCallum 1992). Although both are biologically compelling, decades of research have shown that most forms of summarizing genetic observation are incapable of differentiating these two modes of selection (Hughes and Nei 1989; Meyer and Thomson 2001; Spurgin and Richardson 2010), and the functional insights for the action of heterozygote advantage at least partially explain why it is usually favored over negative frequency dependence (Richman 2000).

A third model involves selective pressures that are heterogeneous over space and/or time, favoring different alleles in different temporal or geographic compartments, and thus resulting in an overall increase in diversity at MHC loci. This model has been shown to be capable of accounting for features of HLA variation (Hedrick 2002). Many studies have investigated this model by comparing the degree of population differentiation at MHC and putatively neutral loci, with the expectation being that selection that is geographically heterogeneous will result in increased differentiation at HLA genes. As reviewed in Spurgin and Richardson (2010), the results are mixed, and interpretation is hampered due to differences in the mutational models underlying the evolution of HLA genes and loci used as neutral controls. Although the specific form of selection acting on MHC genes remains an open question, the fact that these genes have evolved in a non-neutral way and are under balancing selection is an undisputed finding, which is robust to complications introduced by demographic history (Harris and Meyer 2006; Hughes and Yeager 1998; Garrigan and Hedrick 2003).

While studies of MHC have documented convincingly a role of selection, certain questions remain unresolved in the context of variation of the human MHC genes (termed HLA loci). The first of these concerns the “timescale” of selection: while most tests for selection have provided strong evidence for selection at classical HLA class I genes in in deep timescales, there is comparatively less support for selection at recent timescales (Garrigan and Hedrick 2003). It has proved difficult to tease apart the possibility that selection differs across timescales from reduced statistical power of tests for recent selection, and thus the question of the timescale of selection on HLA genes remains open.

The second question concerns targets of selection, i.e, which biological entity is targeted by selection in HLA class I genes: individual alleles or groups of similar alleles? Classical MHC genes have many alleles, which can be hierarchically classified into groups of alleles which reflect the phylogenetic relatedness and shared functional attributes of these alleles. Wakeland et al (1990) proposed a mechanism coined “divergent allele advantage”, which is a specific case of heterozygote advantage, according to which the fitness values of heterozygotes are proportional to the degree of divergence between the alleles they carry. This model was motivated by the observation that, in MHC class II murine genes, alleles from a given allelic lineage often differ by only minor structural variations in the ARS, while alleles in different lineages have functionally different ARS. The open question is whether individual alleles or allelic lineages are the main targets of selection for HLA genes. Although nucleotide diversity intra-lineages exceeds genome-wide averages, inter-lineage diversity is substantially higher than intra (Takahata and Satta 1998). This raises the question of whether intra-lineage variation is under a different mode and intensity of selection with respect to differences between lineages.

We address these questions by analysing the temporal and phylogenetic dynamics of *dN/dS (o*r *ω*) for ARS codons at the class I classical loci (*HLA-A*, -*B* and -*C*) loci, using both pairwise and phylogenetic approaches. These loci are all highly polymorphic and there is an abundance of data available for most exons of their coding sequence, which makes our analyses of non-ARS codons (as a control) possible. Our pairwise comparisons of alleles show that more divergent pairs show higher *ω* for ARS codons than closely related pairs of alleles. The phylogenetic analyses support the hypothesis that selection is stronger for inter-lineage branches (i.e, those connecting two clades from the same lineage, as opposed to those who do not), and also which are internal to the phylogeny (when compared to terminal branches), provided that a bias toward overestimating *ω* for recent divergence is taken into account (Rocha et al 2006). Although evidence for balancing selection on the intra-lineage scale is weaker than on the inter-lineage scale, our findings show that there is statistical support for deviation from a regime of neutrality for intra-lineage branches of the allelic tree. We conclude that intra-lineage divergence has also evolved under a regime of balancing selection, and that inter-lineage divergence bears an even stronger signature of selection.

## 2 Materials and Methods

### 2.1 Data

Alignments for *HLA-A*, *HLA-B* and *HLA-C* were obtained from the IMGT/HLA Database (Robinson et al 2013). All *dN/dS* estimates and related analyses were implemented in CODEML (PAML package, Yang 2007). First codon position was considered to be the first codon of exon 2, as indicated by annotation on IMGT alignments. Our initial data sets were comprised of complete coding sequences, i.e, exons 2-7 (for *HLA-A* and *HLA-C*) and 2-6 (*HLA-B*). These data sets were used for the site models (SM) approach. For the pairwise and branch model (BM) approaches, we used two datasets: one with 48 ARS codons (Chelvanayagam 1996) and the other, referred to as “non-ARS”, consisting of the remaining codons (Table 1).

**Table 1.** Number of alleles and codons for different data sets. a, included all available alleles in release 3.1.0, 2010-07-15., including possible recombinants; b, SM, data set used for site models, i.e, after selection of alleles with complete coding sequences; c, R/NR, with and without recombinants data sets; d, BM (branch models) pruned data set is the NR data set after prunning for alleles which do not cluster intra their respective allelic lineages (see Methods)

| Locus | All alleles<sup>a</sup> | SM (R/NR)<sup>b</sup> | Pairwise (R/NR)<sup>c</sup> | BM pruned data set<sup>d</sup> | Codons |  |  |
| --- | --- | --- | --- | --- | --- | --- | --- |
|  |  |  |  |  | Total | Non-ARS | ARS |
| *HLA-A* | 1193 | 144/107 | 138/104 | 93 | 340 | 292 | 48 |
| *HLA-B* | 1799 | 233/78 | 173/71 | 63 | 324 | 276 | 48 |
| *HLA-C* | 829 | 133/109 | 125/110 | 105 | 341 | 293 | 48 |

In order to be able to use the methods available in CODEML we restricted our analysis to HLA alleles which had complete coding sequences, no stop codons, were expressed in the cell surface and only differed with respect to others by base changes (i.e. no insertions or deletions). Alleles with mutations putatively linked to low or absent cell surface expression were also remove from analyses. The non-ARS data sets were used for estimation of *dS*, used in the pairwise approach as a proxy for allelic divergence, as an internal control for ARS analyses. For the branch models, further pruning of the phylogenetic trees was done, as described below.

### 2.2 Trees and intragenic recombination detection

#### Phylogenetic trees

Complete alignments, described above, were used to generate NJ trees for each gene (Saitou and Nei 1987). The program NEIGHBOR, from the PHYLIP package (Felsenstein 1989) was used with the F84 method, *k* (transition/transversion ratio) = 2 and empirical base frequencies for the distance matrices obtained in DNADIST (Felsenstein 1989).

#### Recombination detection

Intragenic recombinants were detected by applying RDP3 (Martin et al 2010) to the complete alignments, followed by manual inspection. The RDP3 program combines several non-parametric recombination detection methods in sequence data, and we used 6 independent tests for recombination detection: RDP; Chimaera; Maxchi; GENECONV, BootScan and SiScan for recombination detection (see Martin et al. 2010 and references therein). Window size was adjusted to 100 for BootScan and SiScan, and to 15 for RDP. The number of variable sites per window was adjusted to 35 and 30 for Maxchi and Chimaera, respectively. These sizes were chosen based on a test alignment we provided to the software, in which parental and daughter *HLA-B* sequences were known *a priori*. Based on this training set, we adjusted the parameters as described, and for other parameters default values were used. Since these six tests are mostly independent, and have different strengths, we considered a recombination event to be significant when *p* < 0.05 in at least 3 of the above methods, which means we were somewhat conservative in the removal of recombinants from the datasets. “Trace evidence” cases, i.e, those that bear a signal of recombination but are technically not statistically significant, were kept in the data sets. Following this initial procedure, we visually inspected the filtered alignments for the detection of additional recombinant sequences. This procedure generated tow data sets for each locus, one with recombinants and one without (“recombinant” (R), and “non-recombinant” (NR), respectively, Table 1).

#### Clade Filter

For the branch models, we used *t* (expected number of nucleotide substitutions per codon) matrices obtained in pairwise analyses of the non-recombinant non-ARS data sets as input for NEIGHBOR. The trees were visualized for manual pruning and labeling in Mesquite (v2.75, http://mesquiteproject.org/). We imposed that alleles from a given HLA lineage (as defined by the standard HLA nomenclature, which identifies lineage membership by the first field of an allele’s name) had to group together in a clade, and alleles which did not group in such manner were manually pruned from trees in order to fulfill this “clade membership criterium”. The effect of this filtering on inclusion of alleles is presented in Figure S1 in the Online Resource 1. After pruning of the trees, the corresponding pruned alleles were removed from the NR data sets and these reduced data sets were used for the branch model analyses. Table 1 shows the number of alleles used for each analysis.

### 2.3 CODEML analyses

#### Branch models (BM)

With the pruned data sets we compared branch models 0 (one *ω* for all branches) and 2 (two or more categories of branches with independent *ω*) from CODEML. We provided CODEML with a topology based on the non-ARS pruned data set, using branch lengths as starting points for ML estimation (fix_blength=1). For all CODEML analyses (BM, site models and pairwise), the Goldman and Yang (1994) model was used for estimation of substitution rates. Other parameters defined in the control file were as follows: option F3x4 for codon frequency estimation, *κ* = 2 and *ω* = 0.4 as initial values. Tables S14-S16 (Online Resource 1) show likelihood convergence for the branch models, assuming different initial parameter values and codon frequency estimation methods. BM analyses were performed solely for the NR data sets (see tables 2 and 3). Branch models 0 (one omega for all branches) and 2 (two or more omegas) were compared, where branches were labeled either as “intra” or “inter” lineages (Figure 3), or as “terminal” or “internal”. The two models were compared *via* a likelihood ratio test (LRT) with one degree of freedom (see below). BM analyses were performed only for the NR (and pruned) datasets. See Figure 3 for an schema of the labels applied to the trees used in the BM analyses.

**Table 2.** Branch model *dN/dS* estimations and LRT results (ARS data sets). * significance at 5%; Data sets after removal of recombinants (NR); a, *ω* estimate under model 0 (one for all branches); b, *ω* inter lineages; c, *ω* intra lineages d, negative log-likelihood difference between two nested models; e, *ω* for internal branches; f, *ω* for terminal branches

| Locus | $\omega^a$ | $\omega_{\text{inter}}^b$ | $\omega_{\text{intra}}^c$ | $2\Delta l^d$ | $\omega_{\text{internal}}^e$ | $\omega_{\text{terminal}}^f$ | $2\Delta l'$ |
| --- | --- | --- | --- | --- | --- | --- | --- |
| *HLA-A* | 1.84 | 2.03 | 1.68 | 0.06 | 2.35 | 1.39 | 0.49 |
| *HLA-B* | 0.99 | 1.16 | 0.73 | 0.71 | 1.2 | 0.69 | 0.97 |
| *HLA-C* | 1.89 | 4.14 | 1.19 | 2.61 | 4.91 | 0.95 | 7.36* |

**Table 3.** Branch model *dN/dS* estimations and LRT results (non-ARS data set). * significance at 5%; Data sets after removal of recombinants (NR); a, *ω* estimate under model 0 (one for all branches); b, *ω* inter lineages; c, *ω* intra lineages; d, negative log-likelihood difference between two nested models; e, *ω* for internal branches; f, *ω* for terminal branches

| Locus | $\omega^a$ | $\omega_{\text{inter}}^b$ | $\omega_{\text{intra}}^c$ | $2\Delta l^d$ | $\omega_{\text{int}}^e$ | $\omega_{\text{ter}}^f$ | $2\Delta l'$ |
| --- | --- | --- | --- | --- | --- | --- | --- |
| *HLA-A* | 0.53 | 0.40 | 0.77 | 2.8 | 0.39 | 0.95 | 4.57* |
| *HLA-B* | 0.42 | 0.40 | 0.55 | 0.34 | 0.39 | 0.66 | 0.86 |
| *HLA-C* | 0.50 | 0.39 | 0.79 | 3.97* | 0.38 | 0.92 | 5.27* |

#### Site models (SM)

For the SM approach, the clade filter was not applied, which resulted in minor differences between this data set and the other two (pairwise and branch models approach, see Table 1). We used the site models from CODEML to identify codons with *ω >* 1 and thus to test if ARS codons bear evidence for adaptive evolution. M0 (one ratio) assumes the existence of only one *ω* ratio for all codons, while M1 (neutral) assumes the existence of two categories of sites, one with *ω*_1_ = 1 (sites evolving in a neutral fashion) and the other with ω_0_ < 1 (sites evolving under purifying selection), while M2 (selection) adds an extra category to M1, where ω_2_ *>* 1, corresponding to sites with evidence for adaptive evolution. M7 (beta) is a flexible null model where the value is sampled from a beta distribution, where ω_0_ *<* 1, and 0 < ω *<* 1, while M8 adds an extra category to M7, *ω*_2_, which is estimated from the data (Yang 2006). Codons with posterior probabilities P *>* 0.95 of ω *>* 1 in the Bayes Empirical Bayes (BEB) (Yang et al 2005) approach implemented in CODEML were considered to have significant evidence for adaptive evolution, following criteria described elsewhere (Yang and Swanson 2002; Yang et al 2005). The ARS codon classification proposed by Bjorkman et al. (1987) is referred to as BJOR, while the “peptide binding environments”, i.e, the amino acid residues in a fixed neighborhood of the peptide binding residues known from crystal structure complexes (which provide a less restrictive description of the antigen binding sites), are referred to as CHEV (Chelvanayagam 1996). Finally, the list of codons in HLA genes with evidence of *ω >* 1 from Yang and Swanson (2002) is referred to as YANG (Figure 1 and Online Resource 1, Table S9). M1 *vs* M2 and M7 *vs* M8 models were compared through a LRT with two degrees of freedom. Tables S3-S8 (Online Resource 1) show likelihoods obtained when altering initial CODEML conditions for the SM analyses. SM analyses were performed for R and NR data sets.

**Fig. 1.**
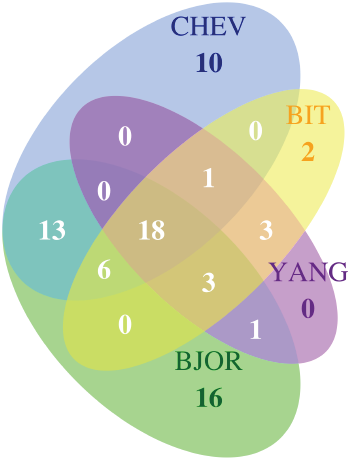
Overlap between two ARS classifications and two site models studies. BJOR and CHEV are ARS classifications (Bjorkman et al 1987; Chelvanayagam 1996); YANG is a list of codons with significant in *HLA* genes; BIT is the set of codons with from our SM (site models) approach (see Materials and Methods for details)

Codons with *P >* 0.95 for *ω >* 1 in M8 (34 in total) were combined for the three loci, and the R and NR data sets, and compared to CHEV, BJOR and YANG. Of these 34 codons, only one was outside of the exons 2 and 3 range (codon 305), which is where all ARS codons are located. Figure 1 shows the overlap between the codons defined as making up the ARS in the BJOR and CHEV classifications, as well as those idenfied as under selection in the YANG set of codons and our analyses.

In order to evaluate if our site model analyses were robust to features of the estimation method, the analyses were repeated with DATAMONKEY, from the HYPHY package (Pond et al, 2005). The substitution model used for construction of the NJ tree was HKY85 (very closely related to F84, used for CODEML analyses). Two criteria for detection significant *dN/dS >* 1 were considered: SLAC and FEL (both with significance level of 0.1), with the former being the most conservative criterion available in the package. Tables S10-S12 report the overlap of sites with evidence for *dN/dS >* 1 for BEB (CODEML), SLAC and FEL.

#### LRT

When comparing two nested models the LRT test statistic is given by doubling the log likelihood differece between the more parameter rich model and the less parameter rich model. The difference in parameter number yields the degrees of freedom. It is expected that the use of a chi-square distribution for significance evaluation of this test is a conservative approach (Yang 2006). Both site models and branch models comparisons were performed through LRTs.

#### Breslow-Day Test

In order to compare ARS and non-ARS codons with respect to the distribution of synonymous and nonsynoymous changes within and between lineages (or for internal or terminal branches), we used a contingency table approach similar to the one described in Templeton (1996). We estimated the synonymous (*S*) and non-nonsynonymous (*N*) changes on each branch in CODEML, using the branch models. Next we counted *N* (nonsynonyous changes) and *S* (synonymous changes) for intra/inter or terminal/internal branches for each locus, and for ARS and non-ARS codons (Table 5).

We defined the odds ratio (OR) as:

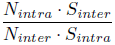

,and used a Breslow-Day test for homogeneity of OR to test the hypothesis that contingency tables from ARS and non-ARS codons have the same OR. We applied the same test to internal/terminal branches. Data from the three loci were combined into the same analysis to increase power.

#### Pairwise approach

We also performed analyses where statistics were estimated in comparisons between all pairs of alleles (pairwise analyses, see Table 1) using runmode=-2 in CODEML. This approach does not require a phylogenetic tree. Because IMGT/HLA nomenclature allows information about allelic lineages to be known without a tree, pairs were also classified as intra or inter-lineage. Correlations between allelic divergence and omega values were tested with a Mantel Test using Pearson’s correlation index (Online Resource 1, Table S13). We obtained quantiles of the *dS*_non–ARS_ distribution and divided pairwise values according to these quantiles (Online Resource 1, Table S1 for non-ARS data set and Table 4 in main text for ARS data set). Differences in mean *ω* values for “intra” and “inter” comparisons were tested for significance by a Wilcoxon rank sum test (Figure 2).

**Table 4.**
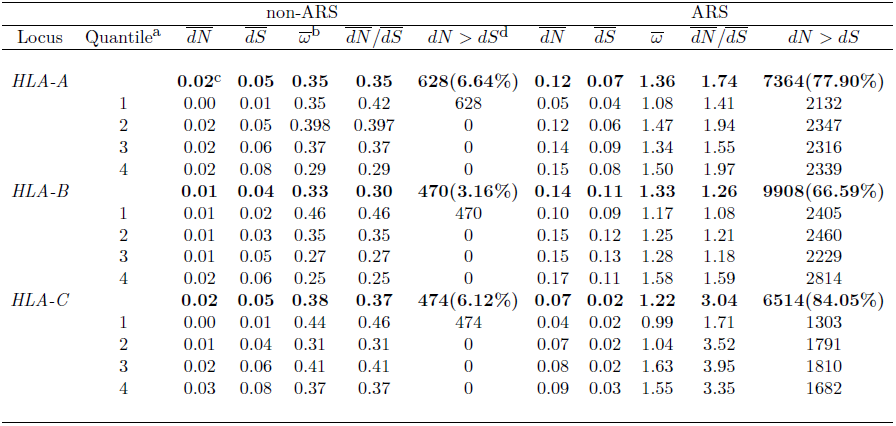
Pairwise estimations for substitution rates (data sets prior to the removal of recombinants). a, quantiles of divergence (*dS*_non-ARS_); b, average pairwise *dN/dS*; c, bold refers to the average pairwise values for each locus; d, percentages correspond to the proportion of pairs for which *dN >dS* in relation to the total number of pairwise comparisons

**Fig. 2.**
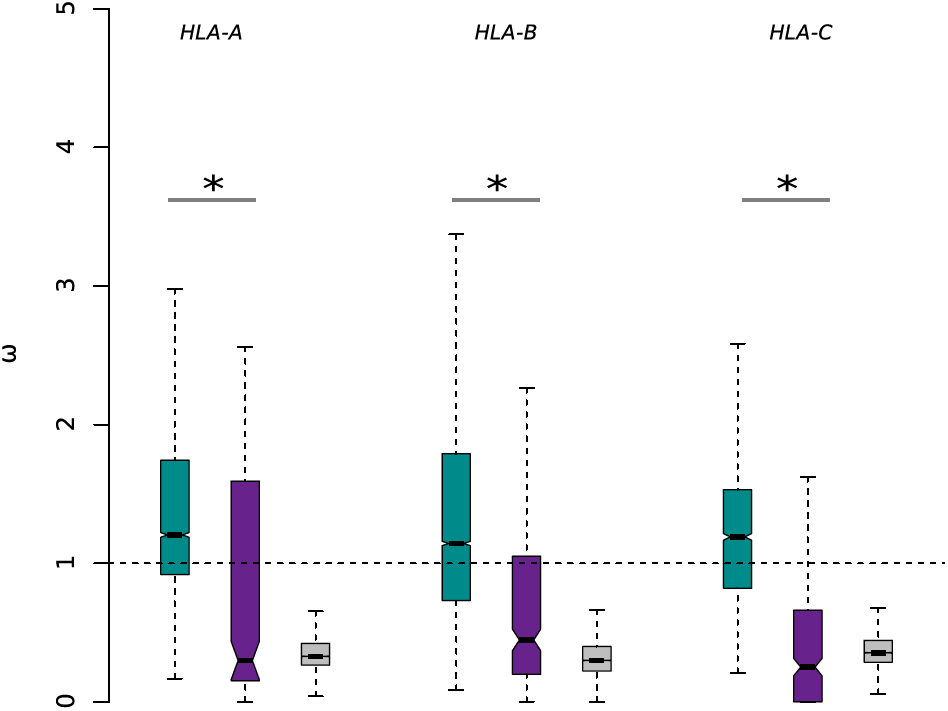
Pairwise estimates for intra-lineage and inter-lineage pairs of alleles. These results refer to ARS data sets prior to the removal of recombinants, for pairwise analyses; Green, inter-lineage; purple, intra-lineage; gray, non-ARS; * significant difference between 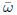 (intra) and 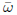 (inter) (*p* < 0.001, Wilcoxon rank sum test)

### 2.4 Allele frequencies of *HLA* SNPs in the 1000 Genomes

The IMGT/HLA database contains all HLA alleles described to date, regardless of their population frequencies. Therefore, it is possible that rare variants can contribute disproportionately to patterns identified in the *dN/dS* analyses. To address this concern, we investigated patterns of variation at the HLA loci in a population (Yoruba, YRI) from the 1000 Genomes Project (1000G), for which frequency of alleles at specific SNP positions is available (*N* = 88 individuals).

To test for a possible enrichment of rare variants in the IMGT data we compared patterns of variation seen in the IMGT and 1000G phase I data (The 1000 Genomes Project Consortium, 2012). To this end, we defined a set of sites, for each locus, which were variable in our IMGT-derived data sets (referred to as the “OVERALL” set of sites). Next, we classified these sites as variable only within a single lineage (“INTRA”), or variable in more than one lineage (“INTER”). For each site, we converted the positions within the HLA locus into a genomic coordinate for *H. sapiens* (hg19).

Next, we verified if these positions are polymorphic in the 1000G Phase I low-coverage dataset (ftp://ftp.1000genomes.ebi.ac.uk) and recorded the minor allele frequency in the YRI population.

## 3 Results

### 3.1 Evidence for selection and assessment of recombination

Before investigating how *ω* varies over time and phylogenetic context, we tested (a) whether selection is detectable in our data set with pairwise comparisons and phylogenetic *dN/dS* approaches; (b) if the presence of *HLA* alleles resulting from intragenic recombination influences our inferences; and (c) if there is agreement between the ARS codons defined by crystal structure (Bjorkman et al 1987; Chelvanayagam 1996) and the codons inferred to have *ω >* 1 in our data set. The results to these tests are pre-requisites for subsequent analyses addressing the more specific hypotheses about heterogeneity in *dN/dS* estimated across the allelic phylogeny and divergence time.

We quantified the mean pairwise 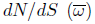, and found 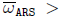 for all loci (Table 4). We used the non-ARS codons from the same sequences as an internal control, and found that 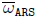 is 3.9 (*HLA-A*), 4.0 (*HLA-B*) and 3.2-fold (*HLA-C*) greater than 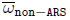 (Table 4). This effect is not driven by a subset of the pairwise comparisons, since *dN >dS* for the majority (between 67 and 84%) of ARS pairwise comparisons, in contrast to the non-ARS comparisons, where fewer than 7% show *dN > dS* (Table 4). Importantly, we find that the result 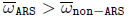 is due to increased 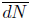 (3.5 to 14-fold higher for ARS), and not to decreased 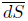 (0.5 to 2.8-fold higher for ARS, Table 4). Qualitatively similar results were obtained when we computed the ratio of mean substitution rates, 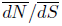 (Table 4). These findings are robust to the presence of recombinants (Online Resource 1, Table S1). Overall, our results document that pairwise comparison of alleles provides strong support for adaptive evolution on ARS codons, as expected.

Evidence for adaptive evolution in ARS codons was also strongly supported by phylogenetic methods from CODEML (see Methods), where models allowing for selection (M2 and M8) in a subset of codons were significantly favored over the neutral models M1 and M7 (Online Resource 1, Table S2; *p* < 0.01, LRT). Results were robust to starting conditions for *HLA-A* and *HLA-B* (Online Resource 1, Tables S3-S6), and less so for *HLA-C* (Online Resource 1, Tables S7 and S8).

We next quantified the overlap between codons we inferred to be under selection (using site models from CODEML, “SM”) and those defined as ARS based on structural analyses of HLA (Chelvanayagam 1996; Bjorkman et al 1987). Within exons 2 and 3 (which contain all ARS codons) we identified 33 codons with significant *ω >* 1 for the M8 site model (see Methods and Table S2, Online Resource 1) in at least one locus, of which 27 (82%) are contained within the set that forms the ARS according to the crystal structure-based classification (Bjorkman et al 1987), 25 (76%) are contained within the peptide binding environments (Chelvanayagam 1996), and 25 (76%) overlap with Yang and Swanson’s (2002) site models approach to detect codons with *ω >* 1 in the three classical class I HLA loci (Figure 1 and Online Resource 1, Table S9). The association between ARS and selected sites for all loci is highly significant (*p < 10^−11^*, chi-square test). There is extensive overlap between the two ARS classifications (Bjorkman et al 1987; Chelvanayagam 1996) (Figure 1) and we also find a high overlap of selected sites between the R and NR data sets for each locus (27 out of 33) (Online Resource 1, Tables S10-S12).

Overall, our results show that: (a) the pairwise and phylogenetic site models methods implemented in CODEML strongly support adaptive evolution on the ARS codons of *HLA* loci – as also described by Yang and Swanson (2002) through site models; (b) there is an enrichment of codons with *ω >* 1 in the CHEV set of codons (see Online Resource 1, Table S9, for the names given to the sets of codons), supporting the use of this classification for our study; (c) although the results were robust to the presence of recombinants, a finding consistent with simulation studies (Anisimova et al, 2003), the estimated values for *ω* appear to be sensitive to the inclusion of recombinants. Therefore, where appropriate, in subsequent pairwise analyses, we contrast results of non-recombinant (NR) and recombinant (R) datasets, while for the branch models we use the NR data set exclusively.

In addition, the results of *HLA-C*, although following the same trend observed for *HLA-A* and *HLA-B*, show that absolute divergence values for ARS codons are on average 1/2 of those observed for the other two loci, both for *dN* and *dS* (Table 4). This result might be a reflect of the fact that *HLA-C* not only has an antigen presentation function, but has a huge role in interactions with NK receptors (KIR) and that, unlike *HLA-A* and *HLA-B*, all *HLA-C* allotypes form ligands for *KIR* receptors (Hilton et al 2015; Single et al 2007). Because the *KIR* loci have been shown to evolve quite rapidly across primate species, plausibly faster than their MHC class I ligands (Single et al, 2007), it is possible that this important selective pressure is responsible for the lower substitution rates seen for the ARS of *HLA-C*, as well as for the lack of consistency observed in ML estimates

### 3.2 The time-dependence of *ω* at HLA class I loci

Having confirmed that selection at ARS sites is detectable with pairwise comparisons and phylogenetic approaches, we investigated if recent evolutionary change (accounting for differences among recently diverged alleles) shows different signatures of selection with respect to changes that occurred over greater timescales. Our first approach consisted in examining the distribution of *ω*_ARS_ as a function of the time since divergence between allele pairs. Our estimate of divergence time between allele pairs was based on the values of *dS* (estimated from non-ARS codons) for each allele pair, thus avoiding statistical non-independence with *ω*_ARS_. Because very recently diverged alleles have low synonymous divergence (*dS*_non−ARS_), the corresponding *ω*_ARS_ values were often undefined or extremely large. We therefore followed a strategy adopted by Wolf et al (2009) to filter out the allele pairs with *ω*_ARS_ *>* 5 (resulting in the removal of 1.1%, 1.4% and 3.9% of *ω* values for pairwise comparisons at *HLA-A*, *-B*, and *-C*, respectively).

Pairwise estimates show that *ω*_ARS_ increases as a function of divergence time (Table 4). Indeed, *ω*_ARS_ and *dS*_non−ARS_ are positively correlated (Online Resource 1, Table S13; *r*_HLA–A_ = 0.17, *p <* 0.001; *r*_HLA−B_ = 0.20, *p <* 0.001; *r*_HLA−C_ = 0.20, *p <* 0.001; Pearson, significance obtained by Mantel Test). Qualitatively similar results were found for NR data sets and were robust to different correlation measures (Online Resource 1, Table S13). We also compared the *ω* between allele pairs classified as intra and inter-lineage (Figure 2). For all loci, the median value of *ω*_ARS_ is *>* 1 for the inter lineage contrasts, and *<* 1 for the intra-lineage contrasts, and the distribution of *ω i*s significantly higher for inter-lineage contrasts (*p <* 0.001, Wilcoxon rank sum test; Figure 2) of the ARS codons.

The above pairwise comparison approach suffers from the limitation that allele pairs with *ω >* 5 were treated as missing data, possibly underestimating *ω* for recently diverged alleles. This prompted us to use a phylogenetic model to contrast alleles at different levels of differentiation, which is more robust to the effects of low differentiation between specific allele pairs. We compared a branch model that estimates a single *ω* for all branches to one that estimates two values of *ω* (inter *versus* intra-lineage; terminal *versus* internal; see Figure 3). For all loci we found higher *ω*_ARS_ for inter-lineage branches than for intra-lineage branches, although significance was not attained for these tests (Table 2). For the contrast between internal and terminal branches, we found higher *ω*_ARS_ for internal branches at all loci and this result was statistically significant for *HLA-C* (Table 2).

**Fig. 3.**
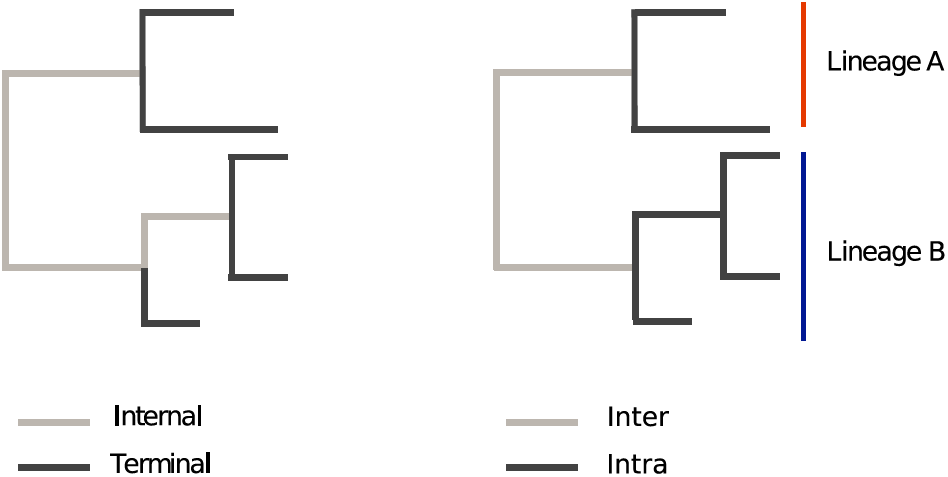
Schematic representation of the allelic phylogenies used in the branch models approach. Left: terminal *vs* internal branches; right: intra-lineage *vs* inter-lineage; For the branch models approach, we labeled branches of each tree (*HLA-A*, *-B* and *-C*) as “intra/inter” or “terminal/internal” and ran model 2 (CODEML), which allows for two independent *ω* values to be estimated, according to these labels

Our results show that both pairwise comparisons and branch models indicate a heterogeneity of *ω* throughout the diversification of HLA alleles, with higher *ω* values associated to contrasts between more divergent alleles (pairwise approach) or to branches connecting different lineages or that are internal to the phylogeny (BM approach), although the difference was not significant for the “intra-inter” contrasts.

### 3.3 Significantly more nonsynonymous changes inter-lineages at ARS codons

In this study we estimate *ω* for allele pairs or branches sampled within a single species, and over varying timescales. Both these features imply in possible biases to the estimation of *ω*, which we now discuss.

Kryazhimskiy and Plotkin (2008) used analytical and simulation approaches to show that under positive selection the behavior of *ω* within a single population is not a monotonic function of the intensity of selection, so that *ω* intra a population can be low, even under positive selection. This occurs because, when an advantageous nonsynonymous variant is fixed in a population, nonsynonymous variation can be decreased due to the homogeneity generated by the selective sweep. However, this scenario clearly does not apply to HLA genes, where balancing selection maintains multiple nonsynonymous polymorphisms simultaneously segregating within a population, contributing to *ω >* 1.

Another challenge to the interpretation of *ω* arises from that fact that many studies have shown that genes under purifying selection show surprisingly high *ω* (often close to 1) when samples with short divergence times are analyzed (e.g., those from a single population or species). For example, Rocha et al (2006) showed that *dN/dS* between two samples is negatively correlated with their divergence times, and exemplified these predictions with bacterial genomes. Likewise, a decrease of *dN/dS* with divergence time has been described in Wolf et al (2009), but considering a much deeper timescale. Kryazhimskiy and Plotkin (2008) demonstrated that this pattern is expected even under a regime of purifying selection that is constant over time. Thus, it is plausible that the recent divergence times among alleles within HLA allelic lineages could result in inflated intra-lineage *ω* values, explaining the modest differences between intra and inter-lineage *ω* values seen in the phylogenetic analyses (Tables 2 and 3). To explore this issue further, we used non-ARS codons as an internal control for this putative build-up of *dN/dS* are recent timescales, and to do so we compared their patterns of variation to those of ARS codons. We found that non-ARS codons have larger intra-lineage *ω* values than inter-lineage values, and also higher *ω* for terminal than internal branches (*p <* 0.05 for *HLA*-*A* in the intra versu*s* inter-lineage contrast, and for *HLA-A* and *HLA-C* in the tips *versus* internal contrast; LRT; Table 3). This distribution of *ω* values is in the exact opposite direction to that observed for the ARS (Table 2), consistent with an effect of short divergence times inflating the estimates of *ω* (Kryazhimskiy and Plotkin 2008).

In order to formally test whether ARS and non-ARS codons have a different distribution of synonymous and nonsynonymous changes intra and inter-lineages (or for internal and terminal branches) we employed a contingency table approach similar to that of Templeton (1996). We used the inferred number of synonymous (*S*) and nonsynonymous (*N*) changes on each branch of the allelic phylogeny from each locus to estimate the total number of each type of change in a specific class of branches (see Figure 3 for a schematic representation of the branch labeling).The odds ratio was defined as presented in the Methods. For all loci, we find that *OR >* 1 for non-ARS codons (proportionally more nonsynonymous on the intra-lineage branches) and *OR <* 1 for ARS codons (proportionally more nonsynonymous changes on the inter-lineage branches), as shown in Table 5. This finding is consistent with the maximum likelihood estimates of *ω* for branches (Tables 2 and 3), and the increased pairwise *ω* inter-lineage, relative to intra-lineage (Figure 2). To test for differences between ARS and non-ARS codons, we pooled the contingency tables of all loci (due to the fact that several cells for individual loci had low counts) and rejected the null hypothesis that contingency tables from ARS and non-ARS codons have the same *OR* (*p − value* = 0.0069; Breslow-Day test). Our analysis comparing internal and terminal branches showed the same pattern, with proportionally more nonsynonymous changes in internal branches for ARS codons (*p − value* = 0.00013; Breslow-Day test; Table 5).

**Table 5.** Distribution of changes for ARS and non-ARS codons. Counts correspond to the total (combined) values for *HLA-A*, *-B* and *-C*; *significant at 1%; *N*, nonsynonymous change; *S*, synonymous change; intra, intra lineage; inter, inter lineage; terminal, terminal branches; internal, internal branches Overall 156 84 17 67

| Data set | Substitution | Branch category |  |  |  |
| --- | --- | --- | --- | --- | --- |
|  |  | intra | inter | terminal | internal |
| non-ARS | *N* | 118.8 | 148 | 89.3 | 158.5 |
|  | *S* | 39.4 | 106 | 24.9 | 115.1 |
|  |  | *OR* = 2.15 |  | *OR* = 2.96 |  |
| ARS | *N* | 172.7 | 230.7 | 144.3 | 291.2 |
|  | *S* | 18.5 | 17.5 | 17.4 | 17.7 |
|  |  | *OR* = 0.71 |  | *OR* = 0.21 |  |
| | | $p = 6.9 \times 10^{-3} *$<sup>b</sup> | | $p = 1.3 \times 10^{-4} *$ | |

In summary, although there is evidence for an excess of inter-lineage nonsynonymous changes (or for terminal branches) for ARS codons, there is also an enrichment for intra-lineage nonsynonymous changes for ARS codons, when compared to non-ARS codons (*P <* 0.001; Fisher’s exact test). Next, we discuss possible biases in the data set which could lead to these results.

### 3.4 Comparing *dN/dS* results with 1000 genomes variation

Our analyses are based on allele sequences available in the IMGT/HLA data base, which is a curated resource to which newly discovered alleles are contributed. This data set is likely to be biased with respect to population frequencies, since very rare HLA alleles are likely to represent a disproportionately larger fraction than in true population samples, since all new alleles which are discovered are encouraged to be submitted to IMGT. We therefore investigated if this bias influenced our findings. Specifically, we were concerned that the enrichment for rare variants could result in an inflation of weakly deleterious nonsynonymous variants for recent divergence, a well documented population genetic signature (Henn et al 2015). This signature could create an artificially inflated value of *ω* for intra-lineage variability.

We found that only a subset of variable positions present in our IMGT-derived datasets are present in the 1000 Genomes Phase I low coverage data (Tables 6, 7, and 8 (for *HLA-A*, *HLA-B*, and *HLA-C*, respectively). This is in accordance with the greater degree of sampling of rare variants in the IMGT data set.

**Table 6.**
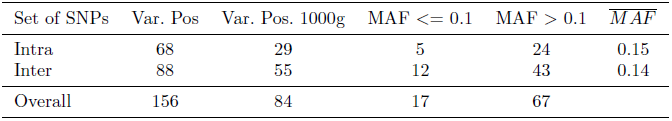
*HLA-A*: MAFs for SNPs in the 1000 Genomes dataset. Overall, set of variable positions considering all sequences in the site models dataset after removal of recombinants. Intra, subset of the ’Overall’ set which is variable only within one allelic lineage for the locus. Inter, subset of the ’Overall’ set which is variable within more than one allelic lineage. Var.Pos, set of all variable positions in the site models dataset. Var.Pos.1000g, subset of Var.Pos which is a SNP in the 1000G low coverage Phase I data. MAF, minor allele frequency. For details, see Methods.

| Set of SNPs | Var. Pos | Var. Pos. 1000g | MAF $\leq 0.1$ | MAF $> 0.1$ | $\overline{MAF}$ |
| --- | --- | --- | --- | --- | --- |
| Intra | 68 | 29 | 5 | 24 | 0.15 |
| Inter | 88 | 55 | 12 | 43 | 0.14 |
| Overall | 156 | 84 | 17 | 67 |  |

**Table 7.** *HLA-B*: MAFs for SNPs in the 1000 Genomes dataset. MAFs for SNPs in the 1000 Genomes dataset. Overall, set of variable positions considering all sequences in the site models dataset after removal of recombinants. Intra, subset of the ’Overall’ set which is variable only within one allelic lineage for the locus. Inter, subset of the ’Overall’ set which is variable within more than one allelic lineage. Var.Pos, set of all variable positions in the site models dataset. Var.Pos.1000g, subset of Var.Pos which is a SNP in the 1000G low coverage Phase I data. MAF, minor allele frequency. For details, see Methods.

| Set of SNPs | Var. Pos | Var. Pos. 1000g | MAF $\leq 0.1$ | MAF $> 0.1$ | $\overline{MAF}$ |
| --- | --- | --- | --- | --- | --- |
| Intra | 44 | 24 | 6 | 18 | 0.30 |
| Inter | 59 | 38 | 8 | 30 | 0.39 |
| Overall | 103 | 62 | 14 | 48 |  |

We next divided positions into two groups: those which are only variable within a single lineage (’INTRA’), and those variable in more than one lineage (’INTER’). For comparison, a third group, which consists of all variable sites (’OVERALL’), was also defined. We found that, considering all INTRA and INTER positions present in the 1000G data, there is no significant difference in minor allele frequency (MAF) between the two categories (Tables 6, 7, and 8). Furthermore, when we classify the 1000G HLA SNPs into low (MAF<=0.1) and high frequency (MAF>0.1), we do not see an enrichment for low frequency variants within the “INTRA” set of SNPs when compared to the “INTER” set (Wilcoxon test, not shown).

**Table 8.** *HLA-C*: MAFs for SNPs in the 1000 Genomes dataset. MAFs for SNPs in the 1000 Genomes dataset. Overall, set of variable positions considering all sequences in the site models dataset after removal of recombinants. Intra, subset of the ’Overall’ set which is variable only within one allelic lineage for the locus. Inter, subset of the ’Overall’ set which is variable within more than one allelic lineage. Var.Pos, set of all variable positions in the site models dataset. Var.Pos.1000g, subset of Var.Pos which is a SNP in the 1000G low coverage Phase I data. MAF, minor allele frequency. For details, see Methods.

| Set of SNPs | Var. Pos | Var. Pos. 1000g | MAF $\leq 0.1$ | MAF $> 0.1$ | $\overline{MAF}$ |
| --- | --- | --- | --- | --- | --- |
| Intra | 78 | 27 | 8 | 19 | 0.26 |
| Inter | 68 | 55 | 19 | 36 | 0.24 |
| Overall | 146 | 82 | 27 | 55 |  |

These results reassure us that the intra-lineage variation we observe is not biased in the direction of extremely rare variants, and that our observation that there is evidence for stronger intra-lineage balancing selection for ARS codons than for neutrally evolving regions (non-ARS) is not a spurious result driven by an enrichment for low-frequency SNPs.

## 4 Discussion

Our study documents a positive correlation between *dN/dS* values and the degree of divergence between allele pairs. This result is supported by phylogenetic analyses, which show higher *ω* values for branches connecting different lineages, or branches which are internal to the phylogeny. A heterogeneous nonsynonymous substitution rate (*dN*) for HLA genes was also reported in a study which found that *dN* for ARS codons is not linearly correlated with divergence time in classical HLA loci (Yasukochi and Satta 2014). By further investigating the temporal dynamics in the *DRB1* gene, these authors showed that this rate heterogeneity is likely the consequence of a reduction in the substitution rates in specific allelic lineages, possibly as a consequence of continuous selective pressure by a specific pathogen. In the present study our goal was to explicitly test for heterogeneity in the *ω* ratios over *a priori* defined groups of alleles (the HLA allelic lineages) and for timescales of divergence (low and high divergence). As was the case with the study of Yasukochi and Satta (2014), we find heterogeneity in the intensity of selection, in our case with evidence of increased selection at deeper timescales than at more recent ones, and for greater selection on inter-lineage branches of the allelic phylogeny, with respect to intra-lineage branches. Our findings indicate that long-term balancing selection has resulted in an enrichment for adaptive changes between allelic lineages for HLA class I genes, with proportionally weaker signatures of molecular adaptation for recent (terminal and intra-lineage branches) than for the inter-lineage and for the internal branches.

Although previous studies have shown that low divergence is often associated to inflated *ω* estimates (Rocha et al, 2006), the phylogenetic analyses carried out in the present work relied on non-ARS codons as a control to show that low divergence times of intra-lineage contrasts does not explain the *ω >* 1 values within lineages, at ARS codons. Thus, while we show that inter-lineage selection is stronger than intra-lineage selection, our results also demonstrate that intra-lineage variation bears a signature of balancing selection.

Recently several papers have drawn attention to the effects of divergence times on *dN/dS* estimation (e.g. Wolf et al 2009; Stolestki and Eyre-Walker 2011), and the complexities of interpreting these values when data is drawn from a single population (Rocha et al 2006; Kryazhimskiy and Plotkin 2008). Our finding of increased *ω*_ARS_ among more divergent alleles (or for inter-lineage branches) is conservative in light of these findings, which predict decreased *ω* for more divergent alleles. We accounted for this effect by using non-ARS codons, which have a similar phylogenetic structure to that of ARS codons (after removal of recombinants) to control for the background inflation of omega in recently diverged alleles, and found that ARS codons have very different distribution of *ω*, with increased inter-lineage evidence for selection, exactly the opposite to what is seen for non-ARS codons.

An important caveat to this interpretation is that the temporal dynamics of *dN/dS* appears to be sensitive to the selective regime which is assumed to be operating. Thus, while several authors have shown that, under purifying selection, increased *dN/dS* at low divergence is expected, positive selection can produce a positive correlation with divergence times (Dos Reis and Yang 2013; Mugal et al 2014), which could account for part of the results we describe in this study. However, the case of directional positive selection, involving the sequential substitution of adaptive mutations, is markedly different from the dynamics of a balanced polymorphism, as is the case for HLA genes.

Assuming that balancing selection has been the main selective regime shaping the molecular evolution of HLA genes, and that heterozygote advantage is one (even if not exclusively) of the mechanisms through which selection has acted upon this system, our finding that inter-lineage *ω*_ARS_ is greater than intra-lineage is consistent with the divergent allele advantage model, according to which heterozygotes for more divergent alleles have higher fitness than those carrying similar alleles (Wakeland et al 1990). Under this model, excess of inter-lineage nonsynonymous changes in HLA genes would be expected, which is a result we have shown for the ARS data set. This model has been shown to explain patterns of variation in the *DRB* locus in Galapagos sea lions, where local allelic divergence at this locus positively influences fitness directly (Lenz et al 2013), and not mere heterozygosity or number of alleles at the MHC locus. Most likely several selective regimes have shaped the evolutionary history of MHC genes, as suggested by previous observations, and our contribution suggests that these selective regimes could be operating alongside with divergent allele advantage.

Our results suggest that groups of functionally related alleles (in our analysis, the allelic lineages) should be regarded as important targets of selection, rather than individual alleles. In line with our observations, it has been proposed that HLA supertypes – groups of alleles sharing chemical properties at the B and F pockets of the ARS region (Sidney et al 1996) – constitute the level of variation that is the primary target of natural selection in *HLA-B* genes (Francisco et al., in press). Since there is a high overlap between allelic lineage and supertype classifications (Sidney et al 1996), our results indicate that attempts to understand how natural selection acts on HLA variation benefit by comparing the effects of selection on the allelic, allelic lineage or supertype levels of variation.

## Electronic Supplementary Material

Supporting tables are available as an additional file.

## Competing Interests

The authors declare that they have no competing interests.

## Author’s Contributions

BDB carried participated in the design of the study, performed analyses, discussed results and drafted the manuscript. RDF performed analyses and discussions. DM conceived of the study, participated in its design and discussion and in the drafting of the manuscript. All authors read and approved the final manuscript.

## Acknowledgements

The authors thank Kelly Nunes for thoughtful comments on the manuscript, Richard Single for comments on the statistical aspects of this work, Aida M. Andrés for general comments and Débora Y.C.Brandt for help with the 1000 Genomes data sets.

This work was supported by the São Paulo Research Foundation (grants #2008/09127-8 to BDB; #08/56502-6 to DM) and Conselho Nacional de Desenvolvimento Científico e Tecnológico (#152676/2011-2 to BDB, #142130/2009-5 to RSF and #308960/2009-2 to DM).Data available in public repositories

https://github.com/bbitarello/dNdS-hla-allelic-lineages

